# Evaluating the diversity of maternal age effects upon neonatal survival across animal species

**DOI:** 10.1101/373340

**Authors:** Edward Ivimey-Cook, Jacob Moorad

## Abstract

Maternal effect senescence is the detrimental effect of increased maternal age on offspring performance. Despite much recent interest given to describing this phenomenon, its origins and distribution across the tree-of-life are poorly understood. We find that age affects neonatal survival in 83 of 90 studies across 51 species, but we observed a puzzling difference between groups of animal species. Amongst wild bird populations, the average effect of age was only −0.7% per standardized unit of increasing age, but maternal effects clearly senesced in laboratory invertebrates (−67.1%) and wild mammals (−57.8%). Comparisons amongst demographic predictions derived from evolutionary theory and conventional demographic models suggest that natural selection has shaped maternal effect senescence in the natural world. These results emphasize both the general importance of maternal age effects and the potential for evolutionary genetics to provide a valuable framework for understanding the diversity of this manifestation of ageing in animal species.

## Introduction

Senescence is commonly described as an age-related physiological deterioration of organismal function typically associated with increasing mortality risk (actuarial senescence) and decreasing fertility (reproductive senescence). Adequately replicated studies report actuarial and reproductive senescence in most species across most taxa (Bonduriansky & Brassil 2002; Descamps *et al.* 2008; Jones *et al.* 2008, 2014; Bouwhuis *et al.* 2009; Waugh *et al.* 2015), with especially well documented senescent declines in natural populations of wild vertebrates (Gaillard *et al.* 1994; Nussey *et al.* 2008a, 2011; Lemaître & Gaillard 2017) and laboratory invertebrates (Rose 1984; Kenyon *et al.* 1993; Bonduriansky *et al.* 2008; Galliot 2012). However, a form of ageing distinct from these manifestations of senescence has also received much recent interest: maternal effect senescence is the detrimental result of a mother’s increasing age on traits associated with an offsprings’ life history or fitness, such as survival, size, growth, and lifespan (Bouwhuis *et al.* 2015; Bitton & Dawson 2017; Clark *et al.* 2017; Lemaître & Gaillard 2017; Lippens *et al.* 2017). While these maternal age effects are attracting increased attention, their distributions across the tree-of-life remain poorly described (Bloch Qazi *et al.* 2017). Thoroughly investigating the prevalence and degree to which these maternal age effects occur will serve to advance our current understanding of trait senescence.

As neonatal survival is profoundly important to longevity and fitness (Crow 1958; Hamilton 1966), this is an obvious focus for demographic and evolutionary exploration of maternal age effects. Demographic models have not yet been applied to data to analyse this phenomenon, but much work has aimed to interpret biological causes the direct effects of actuarial senescence (age-related increases in mortality) by fitting mathematical models to mortality data (Ricklefs & Scheuerlein 2002). The most prominent of such functions used to describe actuarial senescence are the Gompertz, Gompertz-Makeham and Weibull Models (Gompertz 1825; Makeham 1860; Weibull 1951). The Gompertz Model imagines that age-related increases in mortality result from an exponential increase in vulnerability to sources of mortality extrinsic to the organisms. The Gompertz-Makeham Model generalizes this to include an additional parameter to account for sources of age-independent mortality. The Weibull Model views ageing as result of catastrophic intrinsic failure which increases in probability with age and assumes that age-specific causes of death are distinctive, independent and cumulative (Ricklefs & Scheuerlein 2002). While it is debatable whether model fitting can by itself provide insights into the proximate biological causes of ageing, these classical demographic models do provide a convenient method for quantifying ageing rates (Pletcher 1999) especially for the purpose of comparative study (Bronikowski *et al.* 2002, 2011; Sherratt *et al.* 2011). There is no obvious reason for why these same principles cannot be applied to describe age-related maternal effects on neonatal survival.

Several hundreds of models have been proposed to elucidate the proximate causes of ageing (Medvedev 1990), including errors in protein translation, accumulation of free radicals causing cellular damage, damage from heavy metal ions to activation of ageing accelerating mutations, and age–related changes in RNA processing (Harman 1956; Orgel 1970; Eichhorn *et al.* 1979; Medvedev 1986). In contrast, there are few evolutionary models of senescence, and all share the central tenant that senescence is caused ultimately by age-related declines in the efficacy of natural selection (Hamilton 1966). Mutation accumulation (Medawar 1952) and antagonistic pleiotropy (Williams 1957) are evolutionary models that differ in details relating to how genetic architecture constrains the response to selection on age-specific traits. Population genetic models use estimates of vital rates (age-specific survival and reproduction rates) and various assumptions related to gene action to predict patterns of actuarial senescence (e.g. Hughes and Charlesworth 1994), and in particular, population genetic models of mutation accumulation predict Gompertz mortality in adults (Charlesworth 2001). More recently, Moorad and Nussey (2016) applied this approach to quantify how age changes the strength of selection for age-specific maternal effects and to show how these changes cause maternal effects upon neonatal survival to evolve. They predicted that evolved demographic patterns of this manifestation of senescence are qualitatively different from actuarial or reproductive senescence. These differences include possible improvements in neonatal survival with early-life maternal ageing and faster-than-Gompertz declines in neonatal survival with late-life maternal ageing. Furthermore, this evolutionary model ascribes clear and meaningful biological causation to maternal age trajectories in the form of age-related changes in the strength of natural selection. In contrast, the classical demographic models lack clear biological cause.

Moorad and Nussey’s model (hereafter referred to as the Evolutionary Model) derive selection gradients using information relating to demographic structure (age-specific rates of survival and fertility). For this reason, model predictions can be expected to be valid only when populations are near demographic and evolutionary equilibria. As classical demographic models tend not to be justified by evolutionary arguments, we expect that the performance of these models to be relatively insensitive to departures from these equilibria. It is reasonable to expect that natural populations are closer to these conditions than laboratory populations. For these reasons, one test for the predictive value of the Evolutionary Model is to compare its goodness-of-fit to those of classical demographic models and determine if its relative performance improves when fit to natural populations.

In this paper, we address conspicuous gaps in our understanding of maternal effect senescence by performing an extensive systematic review of the literature using meta-analytical methodology. We have chosen neonatal survival as our focus for several reasons: 1) this trait’s relationship to fitness is profound and well-understood conceptually (Hamilton 1966); 2) evolutionary theory explicitly models age-specific maternal effects on this trait (Moorad & Nussey 2016); 3) conventional demographic models of actuarial senescence can be adapted to describe maternal-age trajectories; and 4) associations between the trait and maternal age are observed with sufficient frequency to enable meta-analyses. This study asks two sets of questions about the nature of maternal effect senescence as it manifests on neonatal survival rates:

1. Does maternal age tend to affect neonatal survival in the majority of species across different taxa? Do these effects of age tend to be negative? What features of specific studies appear to predict effect sizes?
2. How well does the Evolutionary Model perform relative to classical demographic models? Does this performance improve in studies of natural population, as we would expect from evolutionary theory?

We find that maternal age effects are widespread across animal species, but maternal effect senescence is a general and important phenomenon in only some groups. The reasons for this variation are as yet unknown and represent an ecological and evolutionary puzzle. However, our demographic analyses provide evidence that natural selection is a causal determinant of this manifestation of ageing, and this represents an important first step to increase our understanding of maternal-age effect variation across species.

## Methods

This meta-analysis followed the Preferred Reporting Items for Systematic Reviews and Meta-Analyses (“PRISMA”) guidelines (Moher *et al.* 2009) (see Fig. 1). A literature search was conducted in July 2017 using the online databases Web of Science and Scopus. Google Scholar was also used, but it failed to produce any papers that were not already duplicated from other databases. Search terms are provided in Supplementary Table S1.

**Fig. 1.**
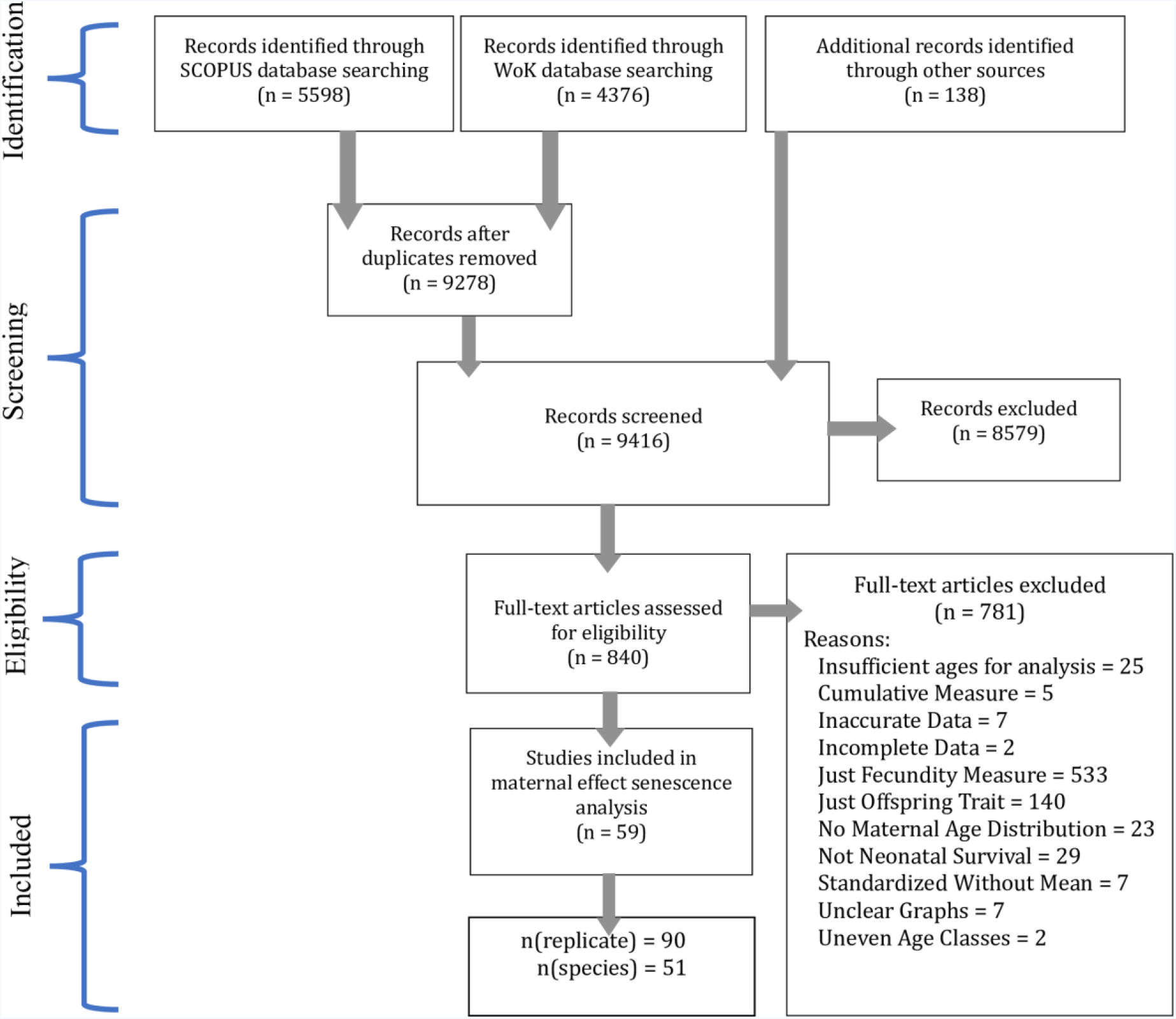
PRISMA flow chart depicting the process and outcome of the literature search.

Accepted papers included the number of surviving and dying neonates as functions of maternal age (see Fig. 1). Papers were rejected if they:

1. had a title or abstract that indicated no appropriate information, or they did not contain data in graphical or tabular forms;
2. couldn’t be accessed;
3. did not contain both fecundity and neonatal survival;
4. focused on humans or highly eusocial animals (as these all have highly complex social systems in which appreciable neonatal care is provided by non-maternal kin);
5. described neonatal survival solely as a function of paternal age; or
6. they included age classes with irregular intervals.

Data were extracted from accepted studies by transcription or by extraction using “WebPlotDigitizer” (Rohatgi 2014), a Google Chrome application that enabled marking of graphical axes, plotting of data points, and conversion to a replicate-specific data file. From each source, we extracted or calculated the following:

1. the number of neonates present at each maternal age class
2. neonatal survival probability at each age class;
3. female age-specific fecundity;
4. cumulative female survival rate;
5. total number of mothers; and
6. the realized maternal probability distribution (i.e. the probability of being a mother at age *x*, calculated as *f*(*x*) = *N_xi_*/Σ*N_xi_* with the *N_xi_* notation representing the number of offspring present at age class *x*).

Binomial datasets were constructed for each replicate in which each standardised age class was associated with a corresponding number of surviving and dying neonates (with corresponding trait values of 1s and 0s, respectively) reconstructed from realised maternal age distribution, age-specific fecundity and neonatal survival rates extracted from the source papers.

We standardized maternal ages by replicate-specific generation time, T, to compare effect sizes across highly variable life histories. For each replicate study i, this was calculated as the average of the maternal age distribution *f*(*x*), or *T_i_* = Σ*_x_xN_xi_*/Σ*_x_N_xi_*. As with any definition of generation time, this measure is sensitive to the age structure and vital rates of the population. This may cause T to change in populations where the timing of breeding is influenced by experimenters who may wish to enhance the power of a study to detect age-related effects rather than to preserve the natural distribution of maternal ages. This likely involves the exaggeration of maternal age variance, and this will tend to increase T compared to natural values. The most likely consequence would be to cause the estimated magnitudes of maternal effects in the laboratory to underestimate those that would be measured in unmanipulated populations.

Studies were identified as belonging to Group N if data came from studies of natural populations, to Group C if data came from semi-captive populations or to Group L if data came from laboratory populations. No species was studied in more than one of these contexts. Classifying studies as describing laboratory and natural populations also effectively separated species into groups with highly disparate phylogeny (Fig. 2) and life histories: bird and mammal species were studied in nature, are long-lived, and provide obvious maternal care; and invertebrate species were studied in the laboratory, are short-lived, and demonstrate little or no conspicuous maternal care. Semi-captive species included vertebrate mammals, birds and reptiles; all provide conspicuous maternal care. More than one binomial datasets were extracted for each species that was studied in different replicates within the same study or in multiple studies. We treated all within-species replicates as independent.

**Fig. 2.**
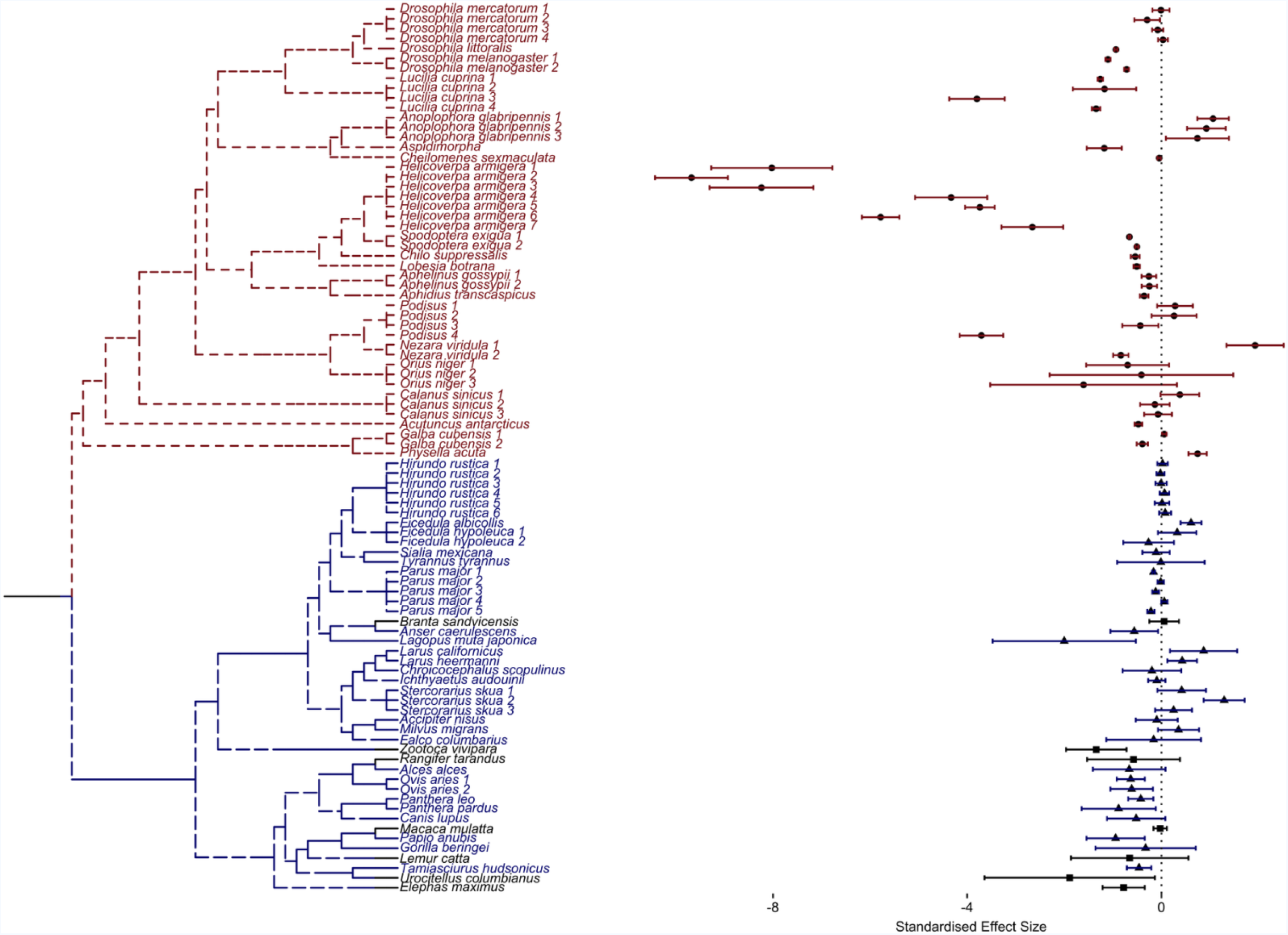
Phylogenetic tree of species included in the comparative analysis with accompanying replicate-specific effect sizes for old maternal age classes (age greater than *T*). The blue text indicates those species evaluated in nature, the red text indicate those assessed in the laboratory, and the black text represents species from semi-captive populations. The forest plot shows the maternal age effect (standardized by generation time, *T*) on neonatal survival across all replicates. Circular points represent effect size estimates for laboratory species, triangular points represent those for natural species and square points represent those for semi-captive species. Error bars around the estimate represent 95% confidence intervals.

Phylogenetic trees were created using the National Centre for Biotechnology Information Taxonomy database (Federhen 2011) (to check taxonomic names for all species) and PhyloT (which converted the list of taxonomic species names into a phylogenetic tree) (Letunic 2011), and visualised using ‘ggplot2’ and ‘ggtree’ (Wickham 2009; Yu *et al.* 2017).

The potential for publication bias should be considered in meta-analyses and tested for statistical significance whenever possible (Egger *et al.* 1997). However, statistical tests were not applicable in this study because those publications that reported maternal age effects quantified these using highly variable methods. For example, some used binomial generalised linear mixed models to report effect size estimates (e.g. Hayward et al. 2015) while other used non-parametric testing with randomisation techniques (e.g. Espie et al. 2000). Some corrected for selective disappearance (e.g. Potti et al. 2013; Hayward et al. 2015), while others did not (e.g. Rockwell et al. 1993; Gagliardi et al. 2007). Quadratic functions of maternal age were fit in some cases (e.g. Newton and Rothery 2002; Blas et al. 2009; Oro et al. 2014); linear functions were fit in others (e.g. Pugesek and Diem 1983; Rockwell et al. 1993). Finally, some studies investigated maternal age effects as only one of many effects of interest (e.g. Baniameri et al. 2005; Jha et al. 2012, 2014), and it may be that publication bias is less likely in these cases as multiple comparisons will increase the likelihood of detecting significant effects.

### Does maternal age affect neonatal survival?

We estimated the effect that maternal age had on the proportion of surviving neonates for each replicate independently. We fit generalised linear models (GLMs) of neonatal survival (*P*) with binomial error (*e*) distribution and “probit” link functions to: [1] age-independent, [2] linear and [3] quadratic models of maternal age (x).

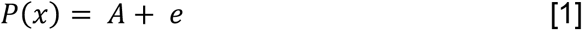

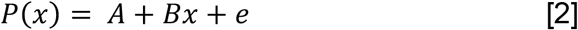

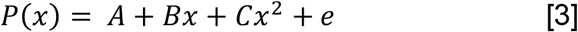

Replicate-specific log-likelihoods for all models were noted along with estimates of effect sizes and associated standard errors. We calculated Akaike Information Criterion values (AIC) for each replicate i, and model j using *AIC_ij_* = 2*k_j_* – 2*loglik_i_*, where *k_j_* is the number of parameters (one, two or three, depending upon the model – see Table 1). From these, sample-size corrected AIC values (AICc) were calculated using the formula 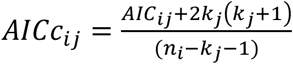, where n_i_ was the number of observations for each replicate (Hurvich & Tsai 1989).

**Table 1.** Demographic models.

| Demographic Model | Survival Function | k |
| --- | --- | --- |
| Gompertz | $P(x) = \exp(-\exp(A + Bx))$ | 2 |
| Gompertz-Makeham | $P(x) = C\exp(-\exp(A + Bx))$ | 3 |
| Weibull | $P(x) = \gamma\exp(-\alpha x^\beta)$ | 3 |
| Evolutionary | $P(x) = \gamma \exp(-\alpha f(x)^{-1})$ | 2 |

Note - The Evolutionary Model predicts neonatal survival based directly upon the reciprocal of the probability distribution function of maternal ages, $f(x)$ (Moorad & Nussey 2016). For convenience, the Evolutionary Model was fit as a Weibull Model but with $\beta$ constrained to be -1 and $f(x)$ substituted for $x$ , but it should be emphasized that this is not a special form of the Weibull function.

### Do maternal age effects tend to be directional?

We used the “boot” package in R Version 3.3.3 (Kushary *et al.* 2000; R Core Team 2016; Canty & Ripley 2017) to calculate the weighted bootstrapped means of maternal age effects estimated from Models 2 (linear) and 3 (linear and quadratic) over all replicates within each species groups (n = 10,000 replicates). Weightings were made by the inverse of the estimated standard errors. Differences between L and N groups were also estimated by weighted bootstrapping.

### Fitting demographic models

Three classical demographic models (Gompertz, Gompertz-Makeham, and Weibull) and a demographic model derived from the Evolutionary Model of maternal effect senescence (Moorad & Nussey 2016) were fit to each replicate (Table 1). All three classical demographic models are intended to describe age-related increases in mortality risk, and these are not sensibly applied to situations where risk declines with age (i.e., increasing neonatal survival with advancing maternal age). The Evolutionary Model allows some initial decline in mortality risk early in life, but it is constrained to predict senescence whenever the maternal age distribution *f*(*x*) decreases with increased age *x*. For every model, we constrained parameters accordingly (see Table S2). Note that Gompertz, Gompertz-Makeham, and Weibull models will converge upon age-independent solutions when neonatal mortality tends to decrease with increasing age, and the Evolutionary Model will converge upon an *f*(*x*)-independent solution when neonatal mortality tends to decrease as selection against neonatal survival decreases. All models were fit as optimisation functions with binomial distributions using the “optimx” package v. 2013.8.7 (Nash & Varadhan 2011) and the “Bound Optimization BY Quadratic Approximation” (BOBYQA) method from the “minqa” package v. 1.2.4 (Powell 2009; Bates *et al.* 2014) and then optimised over two steps in order to increase our confidence that our maximum likelihood solution was evaluated using starting values sampled from a broad range of biologically realistic parameter space:

Step 1: For each of the 90 replicates, 101 models of each demographic model were fit with starting values for intercepts ranging from −1 to 0 (representing neonatal survival that ranged from 0 to 100%) by intervals of 0.01. All other starting parameters were set at 0 or 1 as appropriate. This yielded 9090 solutions for each replicate-by-demographic model family combination. These were then filtered to only include parameter estimates that provided the greatest identified log-likelihood to be used in the next step of model fitting.
Step 2: For each of the 90 replicates, 90 second optimisations were performed using all solutions derived from step 1 as starting conditions. As a consequence of this scheme, initial parameter space for each replicate-by-demographic model family analysis was sampled using reasonable parameter estimates from all replicates. The set of parameters corresponding to the model with the greatest likelihood was judged to be the maximum likelihood solution.

AICc values were estimated using each replicate-by-demographic model log-likelihoods, sample sizes and number of parameters. Calculated AICc values were used to calculate ΔAICc differences and medians between the demographic (Gompertz, Gompertz-Makeham and Weibull) and Evolutionary Models in order to assess overall performance. A different comparative perspective reduced replicate-specific AICc values to a vector of ranks for each model. For example, the model with the lowest AICc is awarded a ‘1’, the model with the second lowest AICc gets a ‘2’, etc. Ranks are summed over all replicates within a species and a new vector of ranks is created from the sum of the component vectors (e.g., the model with the lowest sum of ranks gets a ‘1’). Finally, species-specific rank vectors are summed in the same fashion to obtain species group-specific ranks.

## Results

59 papers met our search criteria. Of these, seven provided data from semi-captive populations (where there was evidence of human intervention in the form of predator exclusion or veterinary intervention), 26 provided data from laboratory populations and 29 derived from natural studies. Some papers included replicate populations (e.g., multiple strains or different environmental conditions for a single species). In total, 90 datasets were extracted and analysed (see Table S3). These replicates represented 20 invertebrate, 13 mammal, 17 bird, and one reptile species. A preliminary search of plant literature was also conducted, however due to low numbers of acceptable papers, we focused our analysis solely on animal species.

### How does maternal age affect neonatal survival?

Replicate-specific results from the GLMs are given in Table S5. As indicated by comparisons of AICc values, the age-independent models were best in 7 cases, linear age effect models were best in 18 cases, and quadratic age effect models were best in 65 cases (out of a total of 90 replicates). Summing AICc values over all replicates indicated a strong preference for the quadratic model of maternal age on neonatal survival (ΔAICc Age-Independent: −81920; ΔAICc Linear: −6721). 69 of the 90 measured offspring outcomes had negative quadratic effects. The weighted bootstrapped means of the quadratic effects were statistically negative when pooled over all species (mean = −0.197, bias corrected 95%-tiles = −0.321, −0.113) and within each group: mean(N) = −0.144 (−0.246, −0.090); mean(L) = −0.212 (−0.414, −0.088); and mean(C) = −0.504 (−1.195, −0.197). The bootstrapped mean difference between L and N suggested that these two groups were not statistically different (mean difference = −0.068, 95%-tiles = −0.109, 0.245). However, the strong tendency across all species towards negative quadratic effects of age indicates that linear models of maternal age tend to underestimate maternal effect senescence experienced by older females (or overestimate maternal effect improvement in the old). In light of this finding, we re-focused our question to evaluate the linear effects of maternal age on old females only, where old defines ages greater than T (i.e., the mothers that are older than average). See Fig S1 for the among-replicate distribution of oldest mothers surveyed.

The distribution of maternal age-effects in old mothers is illustrated in Fig 2. The mean effect of maternal ages was statistically negative over all species pooled together (mean = −0.452, bias corrected 95%-tiles = −0.621, −0.301), over species from Group L (mean = −0.671, bias corrected 95%-tiles = −0.908, −0.456) and over species from Group C (mean = −0.366, bias corrected 95%-tiles = −0.986, −0.073). While the estimated mean effect within Group N was also negative, it was not statistically different from zero (mean = −0.062, bias corrected 95%-tiles = −0.1374, 0.028). As the distribution of effect sizes shown in Fig 2. suggested a profound difference between birds and mammals, we separated Group N into new sub-groups (N_B_ for natural bird studies and N_M_ for natural mammalian studies). In order to test for an overall difference between Groups L, N_B_, and N_M_, we applied a non-parametric Kruskal-Wallis test (n.b. Group C species were removed from this analysis as they were few in number, contained both mammalian and bird species, as well as a reptile, and they exhibited a range of human interventions). We found a significant effect of species grouping on measured late-age effect sizes (χ^2^(2) = 18.399, p <0.001). A Pairwise Test For Multiple Comparisons of Mean Rank Sums (Nemenyi-Tests) from the PMCMR v4.3 package (Pohlert 2014) indicated significant differences between Groups N_B_ and N_M_ (Tukey HSD = 4.625, p = 0.003) and between Groups N_B_ and L (Tukey HSD = 5.415, p < 0.001) but not between Groups N_M_ and L (Tukey HSD = 1.301, p = 0.628). Overall, these results strongly suggest that late-age maternal effects in laboratory invertebrates and wild mammals (N_M_ mean = −0.578, bias corrected 95%-tiles = −0.699, −0.485), are stronger than in natural populations of birds, where the mean effect over all such studies appears to be absent (N_B_ mean = −0.007, bias corrected 95%-tiles =−0.086, 0.085). Note that these effect sizes are scaled as survival fraction changes per generation (e.g., the mean effect pooled over all studies is a 45.2% decrease in survival rates for a +T change in maternal age).

### How well do demographic models fit?

We compared the fits of various demographic models of neonatal survival to extracted data in variety of animal species in natural and laboratory populations. As assessed by median ΔAICc values, the Evolutionary Model performed worse than all three of the classical demographic models (Gompertz: +41.4, Gompertz-Makeham: +26.1, Weibull: +43.9) in laboratory populations (Fig 3A-C). These performance rankings persisted when replicate-specific AICc comparisons were condensed into species-specific model rankings, and ranks were weighted and summed as described above (Table 2).

**Fig. 3.**
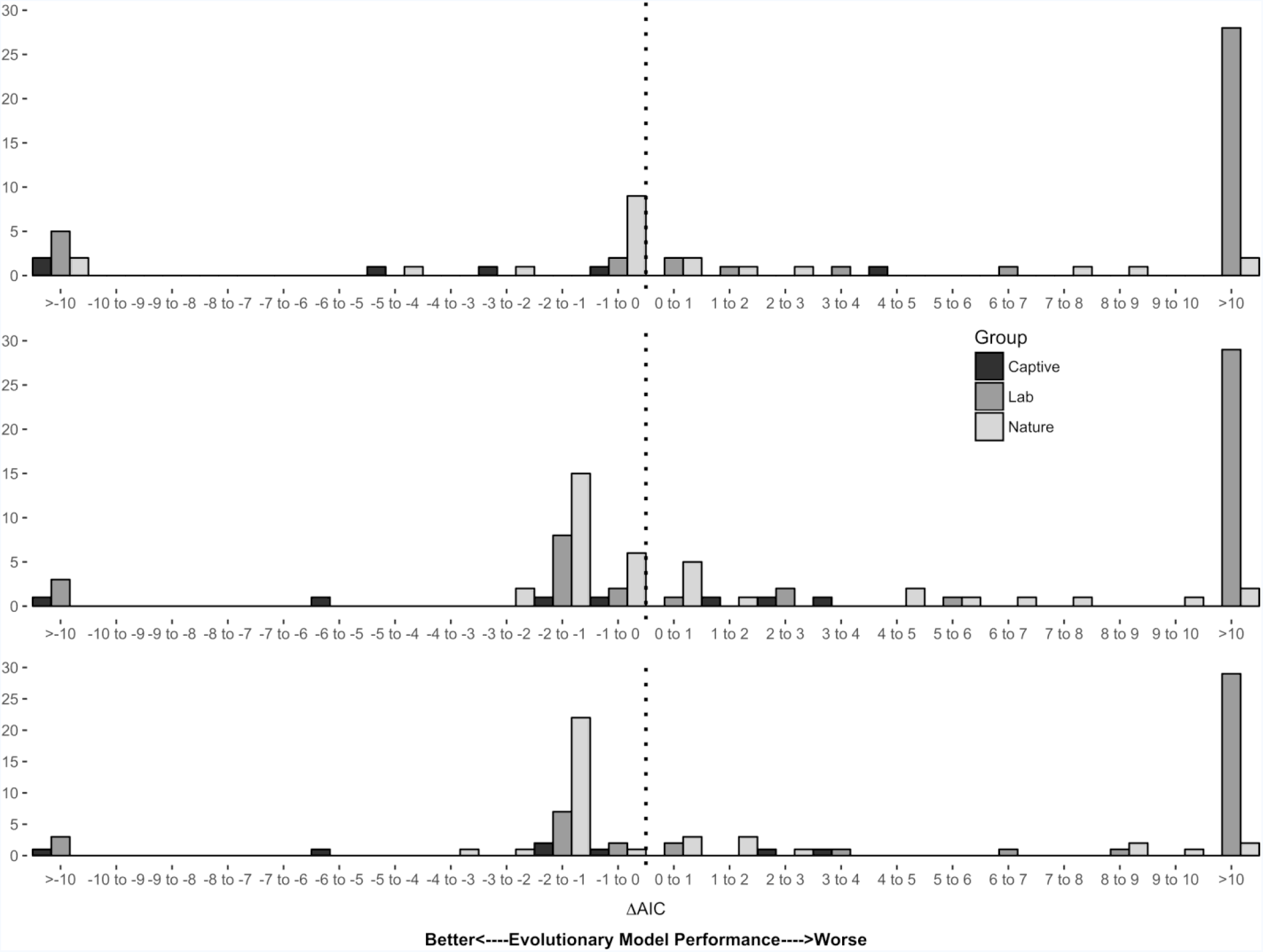
Histograms representing ΔAICc differences between the Evolutionary Model and the A) Gompertz, B) Gompertz-Makeham and C) Weibull models. Bars in black represent counts from captive replicates, dark grey from laboratory replicates and light grey from natural replicates. On 23 occasions, the ΔAICc between the Gompertz and Evolutionary Models was 0; these were omitted from the figure.

**Table 2.** Species group-specific ranking of the four models in Groups L, N and C. Ranked from best (1st) to worst (4th) based on how predictive the four models were when comparing AICcs.

| Ranking | Gompertz |  |  | Gompertz-Makeham |  |  | Weibull |  |  | Evolutionary |  |  |
| --- | --- | --- | --- | --- | --- | --- | --- | --- | --- | --- | --- | --- |
|  | L | C | N | L | C | N | L | C | N | L | C | N |
| 1<sup>st</sup> | 5 | 1.5 | 5.5 | 1 | 1 | 8 | 13 | 1 | 2.5 | 1 | 3.5 | 8 |
| 2<sup>nd</sup> | 8.5 | 1.5 | 9.5 | 7.5 | 2 | 1 | 2 | 2 | 7 | 2 | 1.5 | 6.5 |
| 3<sup>rd</sup> | 4.5 | 2 | 5 | 8.5 | 4 | 9.5 | 3 | 0 | 4 | 4 | 1 | 5.5 |
| 4<sup>th</sup> | 2 | 2 | 4 | 3 | 0 | 5.5 | 2 | 4 | 10.5 | 13 | 1 | 4 |
| Total | 43.5 | 18.5 | 55.5 | 53.5 | 17 | 60.5 | 34 | 21 | 70.5 | 69 | 13.5 | 53.5 |
| Overall Rank | 2 | 3 | 2 | 3 | 2 | 3 | 1 | 4 | 4 | 4 | 1 | 1 |

Note - Lower ranks indicate better models. Scores with 0.5s represent situations in which a tie occurred; an average was taken in these situations.

By comparison of ΔAICc values, the Evolutionary Model appeared to have performed better than the classical demographic models (Fig 3A-C) in natural populations. The median ΔAICc between the Evolutionary and Gompertz Models was −0.242. In 23 cases, the best fit Gompertz and Evolutionary Models both converged on age- and selection-independent solutions with identical log-likelihoods. This lead to identical ΔAICc measures as both types of models share the same number of parameters (two). These non-informative ΔAICcs were removed from the calculation of the median. Gompertz-Makeham and Weibull Models were less favoured in these situations because they fit three parameters; these were included in calculations of these median differences. From median ΔAICc value, the Evolutionary Model outperformed Gompertz-Makeham (−0.941) and Weibull Models (−1.003). When species-specific model ranks were compared rather than median ΔAICc values, the Evolutionary Model performed best (Table 2).

In both comparisons of ΔAICc values and species-specific models ranks in semi-captive populations, the Evolutionary Model was found to outperform the demographic models when assessed by median ΔAICc (Gompertz = −3.565, Gompertz-Makeham = −0.504, Weibull = −1.010) and by summed ranks (Fig 3A-C and Table 2).

## Discussion

### Maternal age effects

Maternal age appeared to affect neonatal survival in 83 of 90 studies accessed in this review (91%), and these effects appeared to be widespread across divergent taxa, life histories and environments. While these results argue persuasively that maternal age effects in late-life are of general importance, phylogenetic constraints may be important in determining whether these effects are directional: increased maternal age clearly tends to become progressively more deleterious in laboratory invertebrates, semi-captive vertebrates, and mammals in nature, but there is no statistical support for widespread late-age maternal senescence in natural populations of bird species. Laboratory populations of invertebrates appeared to experience insignificantly faster maternal-age-related declines in neonatal survival than wild mammals at late ages (67.1% vs 57.8% decline per unit of generation time), but it’s possible that this difference is an underestimate owing to bias associated with estimations of generation time made from experimental studies (see Methods).

On the other hand, there is a clear and dramatic difference between senescence rates between wild mammals and wild birds (57.8% vs 0.7% decline per unit of generation time). This study lacks the means to explain the causes of this difference, but because both sorts of animals provide pre-natal and post-natal care we can reason that it cannot be explained by qualitative differences in the types of maternal care provided. Beyond this, we can only speculate how differences in phylogeny or attendant life-history patterns might generate this variation. One possibility could be that mammalian maternal care is more dependent upon physiological condition than avian maternal care, and this condition degrades with increased maternal age. This borrows from suggestions made in the evolutionary literature that interactions may exist between age-related physiological degradation and condition-dependent environmental hazard that affect age-specific mortality (Williams & Day 2003); this model has been used to suggest that flight reduces that environmental hazard, and this may help explain the oft-made observation that birds live longer than mammals (Williams 1957; Holmes & Austad 1995). As nesting in arboreal or other sites that are protected from predators (such as cliffside or offshore rocks) is often made possible by flight, it could be that flight insulates neonatal birds from effects caused by the physiological senescence of their mothers. Comparisons of maternal age effects on neonatal survival measured in captivity (where the physiological effects of ageing can be suppressed) and in the wild (where they are not) in both bird and mammal species could be made to evaluate this suggestion.

Selective disappearance might explain the dramatic differences between maternal senescence rates in birds and mammals if female survival and maternal quality were more closely associated in birds than in mammals. If so, then female deaths leading up to later ages would cause the preferential removal of poor mothers before neonatal survival could be measured in the post-selection cohort late-in-life; this would lead to a situation in which cohort-level measures of ageing underestimate the true degree of senescence experienced by individuals. Selective disappearance has been discussed at length in the context of actuarial (Vaupel *et al.* 1979; Vaupel & Yashin 1985), reproductive (Bouwhuis *et al.* 2009) and physiological senescence (Nussey *et al.* 2011), but it has only recently been explored in the context of maternal effect senescence (Ivimey-Cook & Moorad 2018). An effect of selective disappearance upon neonatal survival been detected in two studies of seabirds (van de Pol & Verhulst 2006; Zhang *et al.* 2015) but not in a wild population of Soay sheep (Hayward *et al.* 2013) or in a laboratory study of a beetle with conspicuous maternal care (Ivimey-Cook & Moorad 2018), but we lack a biological model that might explain why the phenomenon should be more important in birds than in mammals or invertebrates. Nevertheless, this possibility could be easily evaluated if more future studies of maternal effect senescence report and correct for these effects. This is a simple addition to observational or experimental analyses, requiring only the fitting of time-of-death into standard statistical models, and it should be standard practice for all measurements of maternal effect senescence whenever possible. Finally, we note that the evolution of maternal effect senescence requires an amenable genetic architecture, but, we lack a reasonable biological model that might predict why maternal effect genes in invertebrates and mammals should be more similar in this respect than birds. Perhaps future quantitative genetic analyses (see below) applied to mammal and bird species could shed some light on this possibility.

### Demographic comparisons

Classical demographic models treat age as a predictor of mortality. In contrast, the Evolutionary Model uses age-specific selection, which is derived from age-specific survival and fertility, as a predictor. By both comparative measures used in this study (summed ranks and median ΔAICc values), the Evolutionary Model is superior to the classical demographic models when fit to natural populations. One obvious interpretation is that an age-related relaxation in the strength of selection is a causal determinant of maternal effect senescence, and this manifestation of ageing has an evolutionary component. Added support for this interpretation comes from the relatively poor performance of the Evolutionary Model in laboratory populations, where estimates of current selection should correspond poorly to the long-term intensities of age-specific selection for maternal effects on neonatal survival. However, we cannot ignore the fact that the two environments considered here are not randomly distributed across the tree-of-life; the species represented in wild animal studies as very different from those studies in the laboratory. It is possible that evolution by natural selection has shaped maternal age effects in birds and mammals more than it has caused these effects in invertebrates, but it is difficult to imagine how that difference might have arisen, and this explanations require an effective, but yet-to-be proposed, non-evolutionary mechanism to explain maternal effect senescence in invertebrates. We hope that future work on this subject, such as more studies of maternal effect ageing in insect species with post-natal maternal care (e.g. Ivimey-Cook and Moorad 2018) and observations of senescence in single species assessed in both laboratory and natural conditions (Kawasaki *et al.* 2008).

As with the classical evolutionary theory of senescence (Williams 1957; Hamilton 1966; Charlesworth 1994), the evolutionary model of maternal effect senescence demonstrates that age-attenuated selection is inevitable late-in-life (Moorad & Nussey 2016). However, natural selection can shape evolution only to the degree made available by the underlying genetic architecture (Lande 1979). In the context of maternal effect senescence, this means that: the genetic causes of maternal effects on neonatal survival must be age-dependent to some degree and the ranked-order of these genetic effects must change with maternal age. Direct estimates of maternal genetic effects on neonatal effects in a wild population of red deer (Nussey *et al.* 2008b) provide some evidence that this first condition is met by observing an age-related increase in genetic effects for maternal contributions to offspring birth rate (a predictor of survival). To our knowledge, however, the second evolutionary condition has yet to be tested in the wild. Doing so would involve the measurement genetic correlations for age-specific maternal contributions across ages and testing for correlations that can be significantly bounded away from +1. Such tests should be applied to confirm or refute directly the existence of age-dependent maternal effects on neonatal survival that are inferred by this study.

Finally, it should be emphasized that future conceptual advancements in evolutionary theory could provide better models to explain maternal effect senescence, perhaps by embellishing upon the relative simple population genetic model of Moorad and Nussey (2016). There are many features known to be important to reproductive and actuarial senescence that are not included in this model, such as across-age genetic pleiotropy (Williams 1957; Charlesworth 2001), selective disappearance (Vaupel *et al.* 1979; Vaupel & Yashin 1985; van de Pol & Verhulst 2006), mutational bias (Moorad & Promislow 2008), density- and condition-dependent effects (Abrams 1993; Williams & Day 2003), and within-age trade-offs (Charlesworth & León 1976). In addition to these, cross-generational life history trade-offs or other genetic pleiotropy (e.g. Hadfield 2012) could be important to the evolution of maternal effects. Any or all of these can contribute to extant patterns of maternal ageing.

### Concluding remarks

This study provides the first comprehensive and comparative assessment of maternal age effects on neonatal survival across several diverse animal species. The first goal was to survey across 51 animal species in 59 published papers for interesting distributions of effect sizes; we found that maternal age tends to be an important determinant of neonatal survival across multiple animal taxa. Furthermore, we found that these maternal age effects tended to worsen over time in laboratory invertebrate and wild mammal populations. Surprisingly, this strong signal of senescence was lacking in wild populations of birds. This profound divergence represents a puzzle that deserves future attention. The second goal was to assess these patterns from an evolutionary perspective and to gauge whether natural selection could explain extant patterns of maternal effect senescence. Comparing goodness-of-fits from relevant evolutionary models of senescence to those from demographic models of mortality revealed that the strength of age-specific natural selection was superior to age as a predictor of ageing patterns. Taken together, these findings indicate that maternal age effects upon a trait of fundamental ecological, evolutionary, and demographic importance are widespread and were likely shaped by evolutionary forces.

## Acknowledgements

This research was supported by East of Scotland Bioscience Doctoral Training Partnership (EASTBIO) and the Biotechnology and Biological Sciences Research Council (BBSRC) (544EIC BB/J01446X/1). We are tremendously grateful for the advice given by Ally Phillimore, Jarrod Hadfield, Joshua Moatt and Liam Brierly. We are grateful to members of the Edinburgh burying beetle group for helpful comments on the manuscript. We especially appreciate advice and feedback from Per Smiseth, Alex Scheuerlein, Cammy Beyts, Tom Botterill-James, Jon Richardson, Lucy Ford, Maarit Mäenpää, Matthieu Paquet, and Tom Ratz. The authors declare no conflict of interest.

## Authors’ Contributions

EIC and JM conceived the ideas, designed the methodology, analysed the data and wrote the manuscript. EIC collected the data. Both authors contributed critically to drafts and gave final approval for publication.

